# OrgaQuant: Intestinal Organoid Localization and Quantification Using Deep Convolutional Neural Networks

**DOI:** 10.1101/438473

**Authors:** Timothy Kassis, Victor Hernandez-Gordillo, Ronit Langer, Linda G. Griffith

**Author notes:** **This manuscript is still in draft form and we appreciate reader suggestions before final submission. Please submit comments to ***.

## Abstract

Organoid cultures are proving to be powerful *in vitro* models that closely mimic the cellular constituents of their native tissue. The organoids are typically expanded and cultured in a 3D environment using either naturally derived or synthetic extracellular matrices. Assessing the morphology and growth characteristics of these cultures has been difficult due to the many imaging artifacts that accompany the corresponding images. Unlike single cell cultures, there are no reliable segmentation techniques that allow for the localization and quantification of organoids in their 3D culture environment. Here we describe OrgaQuant, a deep convolutional neural network implementation that can locate and quantify the size distribution of intestinal organoids in brightfield images. OrgaQuant is an end-to-end trained neural network that requires no parameter tweaking, thus it can be fully automated to analyze thousands of images with no user intervention. To develop OrgaQuant we created a unique dataset of manually annotated intestinal organoid images and trained an object detection pipeline using TensorFlow. We have made the dataset, trained model and inference scripts publically available along with detailed usage instructions.

## Introduction

Many of today’s more important biological discoveries have been made using *in vitro* cell culture systems. These systems allow researchers to conduct hypothesis-driven research on a specific cell type to gain mechanistic understanding of its various processes as well as for testing drugs in pharmaceutical research. Conventional *in vitro* cultures have either used primary cells or immortalized cell lines plated on 2D surfaces. While these offer utility, they are not very faithful in recapitulating the complex physiological environment^1^ and are rarely predictive of *in vivo* behavior. Recently there has been a rise in what is called ‘organoid’ cultures^2–5^. Organoids are multicellular spheroids that are derived from either a primary donor or stem cells. In many regards, they resemble their parent organ in both functionality and cellular composition. For example several well received studies have demonstrated the establishment of organoids from the gut^6–8^, pancreas^9–11^, brain^12^, liver^13^ and endometrium^14^ among others. Organoids are fast becoming the ideal model system for understanding development, investigating physiology and for drug testing^3,15,16^.

Obtaining successful organoid cultures that recapitulate the *in vivo* functionality and cellular composition of the target organ requires a tremendous amount of optimization by researchers. Deriving these organoids requires embedding them in biological hydrogels that provide the necessary extracellular microenvironment including growth factors and structural support, and monitoring them over time (days to weeks). Quantifying morphological changes, such as size and shape, as a function of growth or stimulation conditions is fundamental for their use in research. Currently the standard differentiation and culture protocol is to form these organoids inside a gel droplet (**Figure 1a**) that sits on a substrate which is typically the polystyrene bottom of a cell culture multi-well plate or petri dish. To monitor these cultures, the droplet is imaged using a low magnification objective in brightfield (**Figure 1b**).

**Figure 1:**
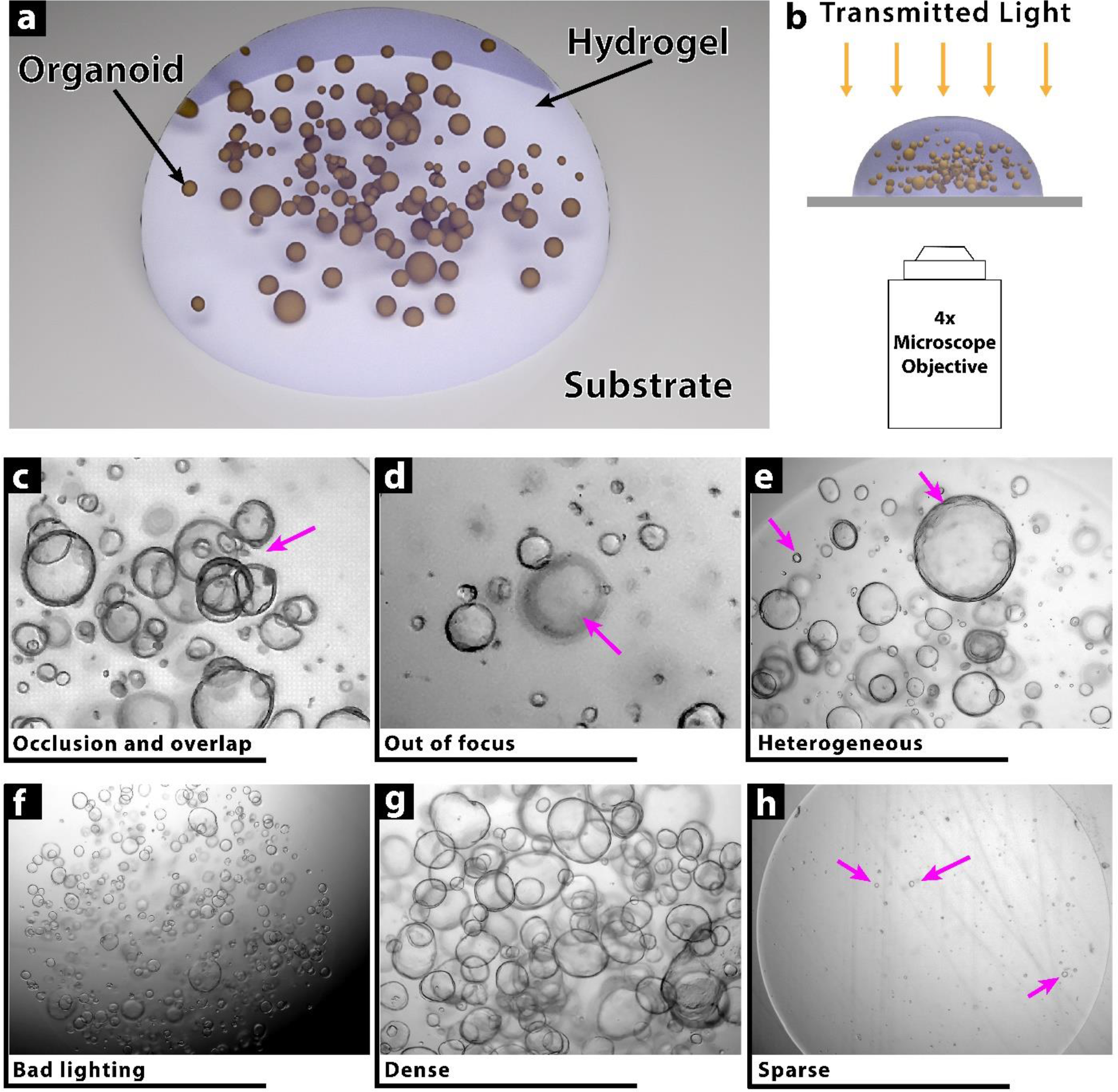
Challenges in Imaging Organoid Cultures. **a)** Organoids are typically cultured in a 3D hydrogel droplet formed from either naturally-derived or synthetic extracellular matrices. The droplet sits on a transparent substrate such as the polystyrene bottom of a multi-well culture plate. Each droplet can have anywhere from zero to several hundred organoids. **b)** Organoid droplets are imaged using low magnification objectives to efficiently capture a wide field of view using brightfield modalities. **c-h)** As a result of the culture and imaging methods there are a variety of imaging artifacts that render conventional segmentation and image processing techniques unreliable. These artifacts include occlusion and overlap **(c)**, out of focus organoids **(d)**, heterogeneous size distribution **(e)**, sub-optimal lighting conditions **(f)**, very dense **(g)** or very sparse **(h)** cultures.

The obtained images typically suffer from numerous imaging artifacts that make conventional image processing techniques extremely difficult. The artifacts include occlusion and overlap, out of focus spheroids, large heterogeneity in size and shape, bad lighting conditions and highly dense or highly sparse organoid distributions (**Figure 1c-h**). Manually measuring and counting these organoids is a very inefficient process as typically there are hundreds of images that need to be quantified with tens to hundreds of organoids per image. As a result, most studies either score by hand a limited number of images or use the images as representative samples and are not quantified.

Recently *Borten et al.* released an elegant open-source software package, OrganoSeg^17^, that addresses some of these challenges, but still relies on conventional image processing techniques and requires tweaking of multiple parameters for any given set of images with similar optical conditions. Instance-based detection using deep convolutional neural networks, however, offers a very promising approach to address this and similar problems. Building on Tensorflow^18^, Google has recently released an object detection API^19^ that makes configuring, training, testing and running various object detection neural architectures substantially easier than before. Utilizing the object detection API, here we present a practical open-source implementation, OrgaQuant, which allows any user to automatically detect and localize an organoid within a typical bright-field image. Based on the idea of transfer learning^20^, we take a pre-trained neural network and further train it on organoid images to achieve very high precision in drawing a bounding box around each organoid. Once a bounding box is determined, downstream processing allows further quantification including size and shape measurements. Using the algorithm does not require any parameter tuning and runs autonomously on all images in a given folder while being robust against the various imaging artifacts described in **Figure 1c-h**. We have made the training dataset, trained model and inference scripts publically available along with detailed usage instructions.

## Results

### A New Bounding-Box Annotated Dataset of Brightfield Intestinal Organoid Images

Since there are no publically available datasets for our model, we created our own unique dataset comprising a total of approximately 14,240 organoids each annotated with bounding box coordinates (**Figure 2**). Please see methods section for dataset-creation workflow. The full dataset including the images and annotations are publically available at https://osf.io/etz8r under an MIT license.

**Figure 2:**
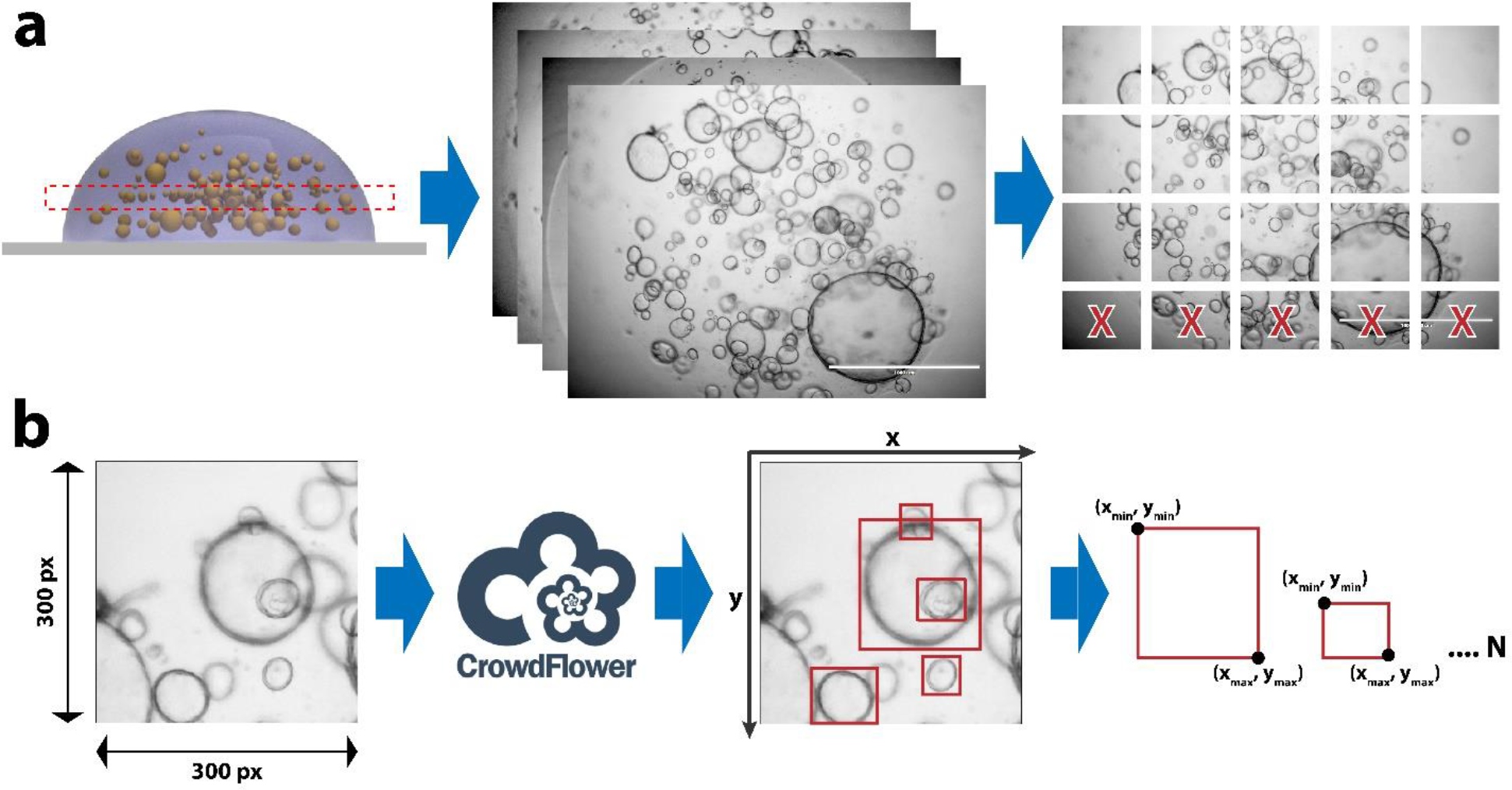
Dataset Creation Workflow. **a)** A single image is captured using a 4× objective which is then divided into 300×300 (and 450×450) pixel patches to make annotations less overwhelming to our crowdsourcing community and to fit within GPU memory of our neural network training computational hardware. **b)** The patches were then distributed on a crowdsourcing platform called CrowdFlower, now knowsn as Figure Eight, along with detailed instructions on what to annotate. Several redundancy techniques were used to assure quality as described in methods. The resulting dataset includes box coordinates (x_min_, y_min_, x_max_, y_max_) for each organoid in an image.

### A Fast, Accurate, Parameterless and Fully Automated Algorithm for Intestinal Organoid Localization and Quantification

OrgaQuant provides a quantification Jupyter Notebook file that can be run to quantify all images within a folder and sub-folders (up to two folder levels deep). The resulting output is a CSV file for each image containing the bounding box coordinates for each organoid, projected 2D area measurements as well as lengths of the major and minor axis of an organoid (which is assumed to be an ellipse). The inference script quantifies an input image by using a sliding window for which both the size and overlap can be set by the user if needed (**Figure 3a**). The sliding window is used to circumvent GPU memory limitations if the entire image where given as input. The output is a single image with all the aggregated labels. Both the labeled image and the CSV labels file are saved in the same folder at the original input image. OrgaQuant labeling quality is indistinguishable from that of humans (**Figure 3b**) with a mean average precision (mAP) of 80%, but is substantially faster and more consistent requiring only 30 sec/patch (on an NVIDIA Quadro P5000 GPU) vs. anywhere from 25 to 284 seconds for humans (**Figure 3c**).

**Figure 3:**
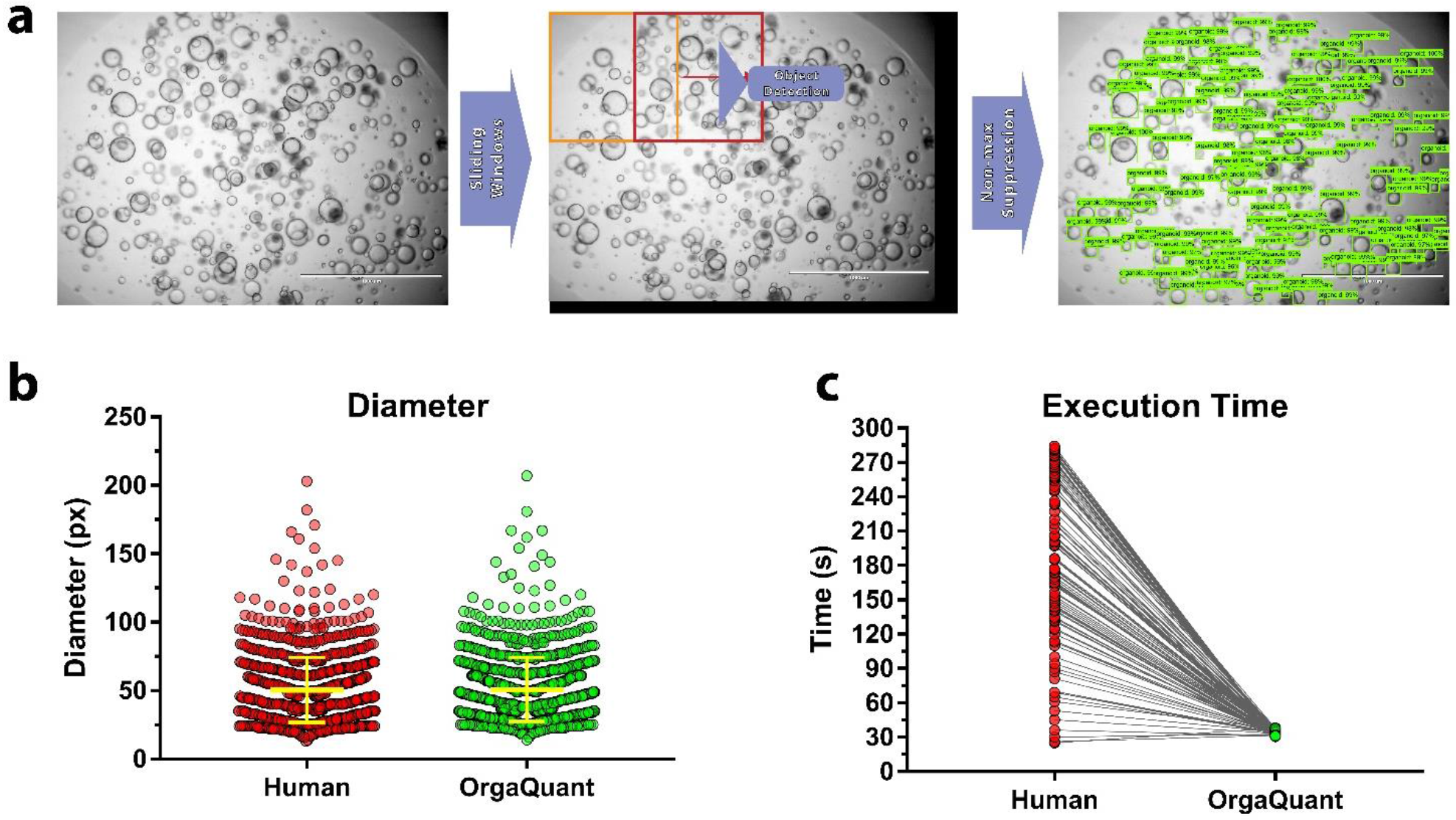
Automated Quantification of Organoids using OrgaQuant. **a)** The inference script runs on individual patches (the size of which can be controlled by the user). To cover an entire image a sliding window with overlap is used. The original input image is padded at the bottom and right edges (shown in black) in order to have an integer number of windows cover each of the horizontal and vertical sliding axes. All the results are automatically aggregated to provide a fully labeled image with the corresponding bounding box coordinates that can be used downstream for any type of processing that might be desirable. **b)** There is no difference between human and OrgaQuant measurements, N= 1135 organoids. **c)** OrgaQuant is substantially faster compared to our trained human annotators and always has the same inference time per patch (around 30 seconds depending on the GPU used). N = 112 different humans.

## Discussion

Object detection and localization is a complex problem in computer vision applications. It is especially difficult when fast detection performance is required. There have been several detection algorithms implemented to provide a balance between speed and accuracy. Two very popular approaches are Single Shot Multibox Detector (SSD)^21^ and You Only Look Once (YOLO)^22^. While these are ideal for real-time detection they accomplish speed by sacrificing accuracy. For OrgaQuant we decided to implement Region Convolutional Neural Network (R-CNN). Specifically what is referred to as Faster R-CNN. Faster R-CNN can use a detection model based on several different architectures including ResNet 101^23^ and Inception v2^24^. Here we chose an architecture based on both Inception v2 and Resent called Inception-ResNet-v2^25^ for which an implementation is provided with the Tensorflow object detection API. The model has been pre-trained on a box annotated COCO dataset^26^ and for our purpose we fine-tuned the model by training it on our organoid dataset. Since Inception-ResNet models have a lot of parameters it is important to use a very large dataset. To achieve this, we augmented the dataset as described in the methods section.

The resulting implementation of OrgaQuant can automatically localize an organoid within a brightfield image and label it with a bounding box. The cropped organoid image can in turn be used in any number of downstream image processing pipelines. Here we demonstrate the ability to measure the organoid size. An important byproduct of convolutional neural networks is that they extract features that can be used with various other machine learning algorithms to, for example, cluster similar organoid based on visual similarily or even detect subtle changes in organoid morphology in response to stimuli that cannot necessarily be detected with normal human vision. Additionally, given the fact that each organoid is localized in 2D space, we can also track the individual growth kinetics of each organoid in a droplet overtime. While we don’t explicitly use OrgaQuant for this, it is as easy as loading a time-lapse set of images in a folder and analyzing it. We believe OrgaQuant is a basis for many exciting and intelligent organoid quantification techniques and we look forward to working the organoid community to further develop this open-source implementation.

## Methods

### Intestinal Organoid Culture

De-identified tissue biopsies were collected from unaffected duodenum areas of children and adult patients undergoing endoscopy for gastrointestinal complaints. Informed consents from the donors’ guardian and developmentally-appropriate approval from donor were obtained at Boston Children’s Hospital. All methods were carried out in accordance with the Institutional Review Board of Boston Children’s Hospital. Tissue was digested in 2 mg/ml of collagenase I for 40 min at 37 °C followed by mechanical dissociation. Isolated crypts were resuspended in growth factor-reduced (GFR) Matrigel (Becton Dickinson) and polymerized at 37 °C. Organoids were grown in organoid expansion medium (OEM) consisting of Advanced DMEM/F12 supplemented with L-WRN conditioned medium (50% vol/vol, ATCC, cat. no. CRL-3276)^8^, glutamax, HEPES, murine epidermal growth factor (EGF, 50 ng/ml), N2 supplement (1X), B27 supplement (1X), human [Leu15]-gastrin I (10 nM), N-acetyl cysteine (1 mM), nicotinamide (10 mM), SB202190 (10 μM), A83-01 (500 nM), and Y-27632 (10 µM) as described^27,28^. Media was changed every two days and organoids were passaged every 4 days by incubating in Cell Recovery Solution for 40 min at 4 °C, followed by trypsin digestion for 5 min at 37 °C to obtain single cells. Single cells were seeded at a density of 25,000 cells in 25 µL of GFR Matrigel. For experiments involving the synthetic hydrogels, single cells were seeded at a density of 500 cells/µL. Three µL of cells suspension (Matrigel or synthetic hydrogels) were loaded in a 96-well plate an allowed to polymerase for 15-20 min at 37 °C. 100 µL of OEM was loaded in each well. Media was changed every two days

### Image Acquisition

Images of organoids suspended in gel droplets were acquired using a Thermo EVOS FL microscope with a 4× objective at day 6 of culture in normal brightfield mode. Images were saved as 8-bit TIFFs along with a scale bar. A single image was taken for a droplet. Since the organoids are suspended in the gel, the focus level was chosen as to have the most organoids in focus as determined subjectively by the user. The resulting images were 1500 × 1125 pixels and were approximately 4.5 MB in size.

### Training Dataset Creation

There are no publically available datasets for labeled organoid images, instead, we created our own (**Figure 2**). Each image (which was around 1500×1125 pixels) was divided into 300×300 pixel and 450×450 pixel patches. It was important to use patches because the original image was 1) too big to fit into GPU memory and 2) too difficult to label as it had hundreds of organoids. The patches were then labeled using a crowdsourcing platform (Crowdflower.com, now known as Figure-Eight) where the workers drew a bounding box around each organoid that was considered to be in focus (i.e. not having very blurry edges). The definition of what ‘in focus’ is very subjective and there was no way to standardize that. Each image was labeled by two different workers and if there was less than 80% agreement (as defined by calculations of IoU carried out by CrowdFlower) the image was presented to a third worker. The bounding boxes that were chosen for each image where an aggregate where a box is only chosen if there was 70% agreement between all workers. Detailed instructions and examples were provided to the workers who could only complete the task after a quality test. Additionally, each labeling task had a test images to assure date integrity. The resulting dataset was composed of 1,750 image patches and a total of 14,242 aggregated bounding boxes. The dataset was divided into a training and test sets. Training had 13,004 boxes and test had 1,135. There were a total of 1,745 unique images that had at least one bounding box. The bounding box data was stored in a ‘.csv’ file where each row contained:

1. **filename**: the image name in which the bounding box is located
2. **width**, **height**: of the image patch (in our case we had two different patch sizes 300×300 and 450×450)
3. **class**: the label for the bounding box. ‘organoid’ was the only label we used.
4. **xmin, ymin, xmax, ymax**: define the coordinates of the bounding box where the origin (0,0) is located in the top left corner of the image.

### Hyperparameter Selection and Neural Network Training

While implementing a Faster R-CNN from scratch is no trivial task. The Tensorflow object detection API made is incredibly easy. While we will not reiterate the steps we took which are well documented on the API’s Github page. We will briefly describe the entire implementation and refer the user to our code for more details.

1. The dataset was created by breaking apart large microscope images of organoids into 300×300 and 450×450 pixel patches.
2. The patches where then uploaded to a Google Storage Bucket to make them accessible to our courdsourced annotators.
3. An instruction manual was written for the crowdsourcing platform called CrowdFlower.com and a new job on the platform was set up to annotate the images using bounding boxes as defined by specific instructions.
4. The resulting ‘.csv’ file included the xmin, ymin, width and height of each bounding box. A small python script was written to change that to xmin, ymin, xmax and ymax as this is the preferred format for the helper scripts used below.
5. The ‘.csv’ file was broken into a training set and a test set.
6. A helper script provided by the API was then used to transform the data from ˙csv format into TFrecords (which is a TensorFlow data format used by the API).
7. A configuration script was then created where we specified the number of classes, augmentation strategy, data location…etc. We also had the option of specifying parameters relating to the Faster RCNN architecture, but we decided to stick with the defaults as that seemed to work well. The hyperparameters we adjusted were:

a. The batch size used was 1 as anything larger did not fit into a single GPU memory.
b. Total training steps for 200k
c. Learning rate was adjusted to decrease with the number of steps as follows:

i. LR = 0.001 from step 0-50k
ii. LR = 0.0001 from step 50-80k
iii. LR = 0.00001 above 80k
8. The training was carried out on a cloud-based Windows Server 2016 instance on Paperspace.com and took around 3 days on a Quadro P5000 GPU with 16 GB of GPU RAM. The service used was Paperspace.com as it was cheaper than both AWS and Google Cloud (for GPU instances).
9. TensorFlow comes with TensorBoard which allowed us to observe the training loss as it was training and to calculate the mean average precision for the implementation (mAP) using the code-base provided by the API.

The main metric we used to evaluate the algorithm’s accuracy was the mean average precision (mAP). This metric is the gold standard for assessing object detection algorithms. The mAP was determined using a 10% held out test set that the training algorithm had not seen. To describe the metric in a bit more detail: The average precision refers to what fraction of the ground truth (manually annotated) bounding boxes were found by the algorithm. For example if an image has two organoids (hence two bounding boxes) and the algorithm detects only one of them, then the average precision is 0.5 or 50%. If it detects both of them then it would be 100%. The mAP is then the mean of all the precisions calculated across all the test images. Hence the closer the mAP to 100% the better is the algorithm. Note that in order to compare the bounding box created by the algorithm with the ground truth, it was assumed if there was 70% overlap (i.e. 0.7 intersection over union) then it was considered the same bounding box. While in some instances it might be useful to have a metric that measures computational efficiency, here it wasn’t a large concern as the implementation did not have to be fast. For example no real-time detection was desired.

### Code and Data Availability

All code for inference is available at https://github.com/TKassis/OrgaQuant, the trained model and full training dataset can be downloaded from https://osf.io/etz8r.

## Acknowledgements

The authors are grateful to Chloe Yang for providing input, to all the contributors to TensorFlow and the Google Object Detection API and to the NIH for funding.

## Author Contributions

Authorship was determined according to a set criteria^29^.

TK: conceived of idea, developed code, ran experiments, analyzed data, wrote and edited manuscript.

VH: ran experiments, analyzed data, wrote and edited manuscript.

RL: developed code and edited manuscript.

LG: provided input, suggestions and edited manuscript.

